# Supporting diamond open access journals. Interest and feasibility of direct funding mechanisms

**DOI:** 10.1101/2023.05.03.539231

**Authors:** Quentin Dufour, David Pontille, Didier Torny

## Abstract

More and more academics and governements consider that the open access model based on Article Processing Charges (APC) is problematic, not only due to the inequalities it generates and reinforces, but also because it has become unsustainable and even opposed to open access values. They consider that scientific publishing based on a model where both authors and readers do not pay – the so-called Diamond, or non-APC model – should be developed and supported. However, beyond the display of such a support on an international scale, the landscape of Diamond journals is rather in the form of loosely connected archipelagos, and not systematically funded. This article explores the practical conditions to implement a direct funding mechanism to such journals, that is reccurent money provided by a funder to support the publication process. Following several recommendations from institutional actors in the open access world, we consider the hypothesis that such a funding would be fostered by research funding organizations (RFOs), which have been essential to the expansion of the APC model, and now show interest in supporting other models. Based on a questionnaire survey sent to more thant 1000 Diamond Open Access journals, this article analyzes their financial needs, as well as their capacity to interact with funders. It is structured around four issues regarding the implementation of a direct funding model : Do Diamond journals really make use of money, and to what end ? Do they need additional money? Are they able to engage monetary transactions? Are they able to meet RFOs’ visibility requirements? We show that a majority of OA Diamond journals could make use of a direct funding mechanism with certain adjustments. We conclude on the challenges that such a financial stream would spur.

## Introduction

More and more academics and governements consider that the open access model based on Article Processing Charges (APC) is problematic, not only due to the inequalities it generates and reinforces (Ellers et al., 2017; Ross-Hellauer et al., 2022; Smith et al., 2021), but also because it has become unsustainable (Khoo, 2019; Morrison et al., 2022) and even opposed to open access values (“The Budapest Open Access Initiative:20th Anniversary Recommendations,” 2022). They consider that scientific publishing based on a model where both authors and readers do not pay should be developed and supported (Becerril, Arianna et al., 2021; Miedema, Frank et al., 2020). But beyond the display of such a support on an international scale, how to concretely achieve it, when the landscape of non-APC journals is rather in the form of loosely connected archipelagos (Bosman et al., 2021)? This paper is a contribution to this ongoing movement that seek to build a post-APC landscape (Estelle and Wise, 2023). It explores the practical conditions to implement a direct funding mechanism to such journals, that is reccurent money provided by a funder to support the publication process. This funding mechanism would be fostered by research funding organizations (RFOs), which have been essential to the expansion of the APC model, and now show interest in exploring and supporting other models (Ancion et al., 2022; Yang et al., 2023).

At first sight, the APC model is the predominant revenue model of open access journals publishing, in which a fixed sum given by authors is the price to publish a given article in open access. Popularized in the 2000s by academic publishers such as BMC and PLOS, this model spread thanks to the massive support of some private and public RFOs that made their cost eligible in grant money. Despite its visibility, the APC model is still being adopted by a minotity of open access journals. It tends to outshine a wide variety of alternatives, currently referred to as “Diamond open access”. Interestingly, this label is often defined negatively as “non-APC journals”, that is, the absence of a requirement for authors to pay to publish in open access. For instance, the Directory of Open Access Journals website displays a “Without APC” filter for journal search. But that encompassing label does not say much about the actual sources of funding and support, the diversity of these journals business models, and their associated costs.

In order to clarify the content of this catch-all label, some authors tried to break down the diversity of models that it embraces, beyond the already long suggestion list written in the Budapest Open Access Initiative two decades ago (BOAI, 2002). For example, Peter Suber (2007) suggested to range them from direct institutional support to advertising model. This literature, which is small in volume, is almost always based on the same idea: offering business models to journals currently under subscription in order for them to flip to open access (Laakso et al., 2016). Most recently, Wise and Estelle articles (2020, 2019) on “Plan S-compatible” transition models to open access for scholarly societies are prototypes for this approach. In addition to the various forms of APC, the authors identify alternative options: “transformative” models, i.e., redirecting library subscription fees to open access publishing; establishing a cooperative publishing infrastructure between the publisher and libraries; allowing self-archiving of “accepted author manuscripts” or postprints; open publishing platforms (F1000, Emerald Open Publishing), i.e. a preprint repository with paid peer review services; “other forms of funding” which include freemium, subsidy, crowdfunding, publication, syndication, “subscribe to open” model; and finally cost reduction (eliminating or combining journals, pooling management software, stopping paper production, etc.). In the same vein, the recent Open Access Diamond Journals study (OADJS) displays some elements about non-APC business models (Bosman et al., 2021). While describing for the first time a widespread ecosystem (nearly 30,000 journals in varied disciplines from SSH to STM), the report underlines the importance of non-directly monetary contributions. It also shows the diversity of monetary sources necessary for their functioning: grants, donations, crowdfunding, shared infrastructures, institutional support model or freemium. Additionally, some studies focus on the business model of third parties that would fund journals, would it be dissemination platforms (Mounier, 2012), libraires OA funds (Verbeke and Mesotten, 2022) and the “subscribe to open” model to transform the relationship between libraires and publishers (Crow et al., 2020).

Our aim is to expand the Diamond OA journals revenue models beyond these collective funding initatives (Pooley, 2021), by focusing on important actors of the research ecosystem, namely RFOs, usually absent from this literature. While RFOs are still central in the APC world that they helped to expand, notably in the health sciences (Solomon and Björk, 2012), they appear to be vastly underrepresented in the Diamond OA world, with only 5% of journals mentioning them (Bosman et al., 2021)^1^. Furthermore, when Diamond OA journals were asked in an open-ended question about their ideal funding model, only 4% mentioned RFOs as a possible or desirable source^2^. However, when RFOs were directly mentioned in a question about the help expected by journals, they largely responded, ascribing to them the provision of services, grants and donations, but also structural funding (Bosman et al., 2021, p. 98).

Our article is a followup of the OADJS recommandations (Becerril et al., 2021) in which one of us took part (see Appendix 1). We explore the conditions of implementation of a key recommendation stated in the OADJS report, namely a direct funding mechanism for Diamond OA journals. Such a mechanism would give RFOs a permanent funding role in the Diamond ecosystem. If RFOs contributed to create and maintain the APC model by financing it, we assume it is possible for them to reallocate financial flows to expand the Diamond OA journals ecosystem.

The first section of the article is dedicated to the methodology we followed, notably based on the OADJS ressources (1). We then tackle a variety of questions to adress the conditions of implementation of a direct funding model. The second part refers to the actual monetary needs of OA Diamond Journals (2). Do they really make use of money, and to what end ? In the third section, we explore the additional financial needs of such journals, that is, money they would make use of if they where in a hypothetical world with unlimited funds – what we call the “ideal world” (3). To make use of money, OA Diamond journals need to be able to engage monetary transaction: we discuss this point in the fourth part (4). Finally, we explore OA Diamond journals’ ablity to meet RFOs requirements as a counterpart for receiving direct fundings (5).

### 1. Methods

To explore the conditions of implementation of a direct funding mechanism for Diamond OA journals, we developed a questionnaire survey between March and June 2021. In the first part of this section, we give some insights into the elaboration of the questionnaire. We then focus on a specific part of the questionnaire, that is, a list of publication acts that take part in the publication process. We subsequently underpin how we disseminated the questionnaire and collected data, then describe how we analysed the quantitative and qualitative data.

### 1.1. Building the questionnaire

The questionnaire is structured in five sections (see Appendix). The first one delivers basic information about the journal (e.g. title and ISSN). The second part tackles its economic configuration, that is, its embeddedness into a wider structure (or its independence vis-à-vis other structures). It also addresses questions about the journal’s ability to engage in monetary transactions. The third section refers to the publication process and its associated financial needs. For each of the 26 acts that describe the journey of a manuscript from submission to publication (see below), we ask three questions: who performs the task? Does it involve a monetary transaction? In a world without financial constraints, would you pay for this task? The fourth section considers the relationship of journals with a potential funder, notably the capacity to meet funders’ needs and visibility through reporting practices. Finally, the last section comes down to Diamond OA journals’ opinion concerning a direct funding mechanism and the form it would take. The questionnaire has been thought as an OADJS followup study. Closely connected to the OADJS members, we had the opportunity to discuss with them the first draft of our questionnaire. When stabilized, the questionnaire was uploaded to Survey Monkey, an online software program that helps to develop the questionnaire, to send them to the respondents, but also to collect and analyse data.

#### 1.2. A list of acts to describe the publication process

To decompose the publication process in a list of acts, we began by mapping the range of tasks that OA journals must handle in order to go from manuscripts submission to published articles. The aim was to specify the type of resources mobilized for each act, and thus to identify the uses of support that a funder might provide. In order to elaborate an extensive list of acts handled by a journal, we relied on a body of literature that includes cost breakdown presentations by publishers (e.g IOP, Copernicus, MDPI…), a text published on The Scholarly Kitchen website (Anderson, 2018), and several academic works on publication costs (Contat and Gremillet, 2015; Grossmann and Brembs, 2021; Waidlein et al., 2021).

Our exploration of the literature led to the elaboration of a list (table 1), each item corresponding to a single act, providing a rather linear, step-by-step representation of a text’s publication process within an academic journal. The respondents had access to this list of 26 acts, ranging from the receipt of the submitted manuscript to its dissemination as a published document, through the evaluation and the physical production of the format (e.g. scientific article, editorial, proceedings…).

**Table 1.** A list of 26 publication acts

| Name of the publication act | # |
| --- | --- |
| Receipt of a manuscript | 1 |
| Formatting the manuscript before entering the editorial process | 2 |
| Communication with authors | 3 |
| Finding reviewers and monitoring their work, schedule... | 4 |
| Reviewing (definition of criteria, written evaluation format, traceability of exchanges) | 5 |
| Decision of the editorial board (procedures and archiving) | 6 |
| Response to the author (acceptance, rejection, revision) and management of the re-submission process | 7 |
| Plagiarism check | 8 |
| Identification of the author(s) | 9 |
| Conflict of interest check (declaration form and archiving) | 10 |
| Copy-editing (grammar, spelling, style...) | 11 |
| Language editing | 12 |
| Checking compliance with the template | 13 |
| Graphic work (figures, graphs, tables, photos, transcription conventions, etc.) | 14 |
| Proofreading and checking the integration of changes | 15 |
| Translation (summary, keywords, full article...) | 16 |
| Format production (PDF, HTML, XML) | 17 |
| Semantisation of references | 18 |
| Image rights | 19 |
| Managing licences | 20 |
| Rights management and author's contract | 21 |
| Addition of metadata | 22 |
| Assigning a DOI | 23 |
| Integration of the manuscript into an issue | 24 |
| Putting the document online and making it accessible | 25 |
| Publication of data associated with the article | 26 |

Two important precautions must be taken regarding this list of acts. First, its construction obviously tackles theoretical and methodological problems: other categories could have been considered, some refined or clarified, obviously leading to different answers. In order to identify possible gaps in the categories selected, we have associated an open-ended text zone in which the respondents were able to specify elements that did not fit into the formalism of our table. Some acts, such as website maintenance or archiving, were absent from our list as the aim was neither to be exhaustive nor to take into account the singular reality of each journal. With these 26 acts, we strived to grasp the main tasks commonly shared by the journals. Secondly, the number of answers about the publication tasks can vary from one question to the next. This led to a presentation of the final results in rates rather than in absolute numbers of journals. As the number of responses remains high, this part of the survey enables us to draw trends concerning the publication process and its financing.

#### 1.3. Data dissemination and collection

In addition to benefiting from the feedback of the OADJS team, we also relied on their database. The OADJS survey has produced a population of 1,252 journals email addresses that agreed to be contacted anew by members of the team. This population has a significant advantage for our study: since we already had information about these journals (discipline, country, journal age, etc.), we were able to be more specific. Our own survey was opened on June 16, 2021 via an email sent to the 1,252 journals. After two reminders, it was closed on July 12, 2021.

296 respondents opened and started to fill in the questionnaire. After extraction of the data on Excel spreadsheets, we proceeded to their cleaning: deletion of duplicates and uncompleted forms. In the end, we reach a total of 260 journals whose answers were usable for the analysis. Of these 260 journals, 253 could be linked to data produced by the OPERAS survey by matching on ISSN or the name of the journal reported^3^. This population is still very diverse (at least 29 different disciplines and 55 different countries of activity) but the matching allows us to state that the sample of our respondents is close to the OADJS population as the distribution by major groups of disciplines (HSS, Science, Medicine, Multidisciplinary) and the distribution by geographical areas are very similar. The sample of journals is thus dominated by Western Europe countries and HSS journals, but leaves room for great geographical and disciplinary diversity among a half of the respondents.

The number of answers to the different questions varies between 200 and 260 for the general questions, goes down to less than 70 respondents for conditional questions, and to only a few of them for subsidiary questions. When we state that 260 responses are usable for the analysis, we must keep in mind the diversity of respondents’ roles who answered on behalf of their journal. The status within the journal (editor-in-chief, editorial board member, editorial secretary, etc.), the degree of knowledge of the publication process, as well as the forms of investment in the work of publication, may vary from one respondent to another. Answers to the open-ended questions attest to this diversity, since they range from extremely detailed comments to vague or even non-existent comments.

#### 1.4. Quantitative analysis

The majority of the questionnaire was structured by closed questions with predefined answers. Thus, many of the data was processed quantitatively and graphically visualized. For most of the quantitative treatments, we took the dataset for a question, built a table compiling the information, and produced a graphical representation. Since the number of answers from one question to another vary, especially concerning the list of publication acts, we systematically displayed rates rather than absolute values. We also performed some cross-referencing of data sets generated by the answers to several questions. Here again, we made use of the OADJS to put our results in perspective.

#### 1.5. Qualitative analysis

Several questions in the questionnaire were open-ended: they called for open text responses that could not be compiled and integrated directly into quantification processes. We distinguished two types of open text questions for which we carried out different types of processing.

The first type refers to questions with a large number of answers (at least 200), such as the last two questions (5.1. and 5.2.), which collect the respondents’ opinion on a direct financing method for Diamond OA journals, but also the questions associated with publication acts (3.4.1., 3.4.2., 3.4.3.). To work on these data, we exported the answers to a text file and used the qualitative processing software ATLAS.ti. The latter allows us to carry out inductive coding as we read, and ultimately to organize the material according to several categories and sub-categories. For example, question 5.2. asked about the Diamond funding models that journals were considering. Coding within this software allowed us to identify several general categories (infrastructure funding model, service provision model, direct funding models, conditions for implementing a given model, source of funds, desired amounts), which were further broken down into several subcategories (e.g., the direct funding model contains advertising, fundraising, voluntary contributions, fixed sum allocations, or volume funding of publications). Once the categories were stabilized and the coding done on the entire textual corpus, it was possible to find all the text associated with a particular code, and thus to quickly collect examples.

The second type of open-ended questions were those for which the number of answers is very low, usually less than 10. We manually processed the data by reading them directly into the Excel spreadsheet to formulate an interpretation. The usefulness of the answer to this type of question varies greatly.

### 2. Do Diamond OA journals use money and to what end?

A major condition to implement a direct funding mechanism for Diamond OA journals relates to their effective financial needs. Do Diamond OA journals, often portrayed as volunteers journals or “zero cost” initiatives, need money to last? How are these needs distributed among the acts of the publication process? Without financial needs, whether based on “zero cost” or full support by the owning or governing institution, the very idea of a funding mechanism is no longer relevant. By contrast, if these financial needs exist, we strived to precisely locate them on the publication process. In this section, we examine the current monetization of publication acts within Diamond OA journals.

Considering our list of 26 acts, after having anwswered who was their main performer, Diamond OA journals declared which acts were subject to monetary transactions. As shown in the figure 1, for a given act, the answers are globally distributed in a binary manner between “yes” for an act subject to payment, and “no” for an act performed without explicit direct financial support. The third possible answer (“I don’t know”) exists in fairly low proportions, up to 5% for the first 15 acts, and between 2% and 7% for the others.

**Figure 1.**
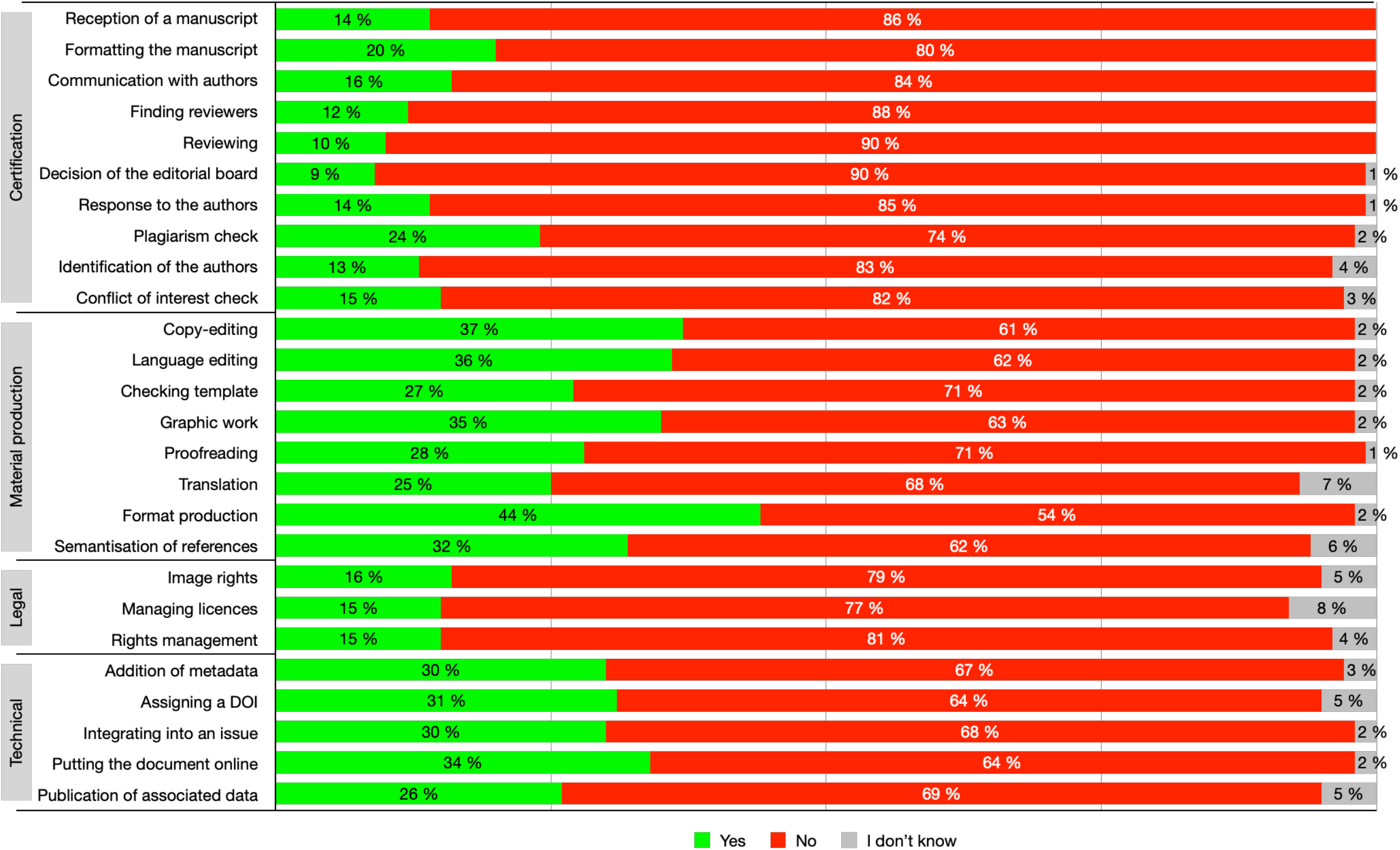
Monetizing publication acts

We emphasize three distinctive results. First, in our sample of Diamond OA journals, the majority of publication acts are carried out without monetary transaction. We are in a situation typical of the general economy of scientific publication, where the work done by academics within journals is an integral part of their activity and is not subcontracted. The publication activity is indirectly remunerated in the salary that institutions pay to persons (i.e. researchers or secretaries) working for the journal, as explained by one of our respondents:

*ID 12778607986: “If I do [the tasks], I do them as part of my general workload and salary as a professor and by making sacrifices elsewhere (usually to my own research work)”*.

This result is in line with the OADJS findings that already highlight this aspect of the Diamond ecosystem.

Second, the publication process is not solely based on such work performed without monetary transaction. Within each journal, the production of published texts involves human and technical resources that are partly based on some monetary exchanges, as the green line shows. Hence, we can state that Diamond OA journals do have monetary needs, that are used to pay in-house personnel, servce providers and subcontractors. Money from research funders could thus be useful if such a flow would be designed.

Third, the monetization of publication acts is distributed between four groups: a first group from act 1 to 10 for which the monetization is low; a second group from act 11 to 18 for which monetization is relatively high; a third group from act 19 to 21 for which monetization is also fairly low ; a fourth group (acts 22 to 26) for which monetization is a bit higher. Such a distribution is an empirical result that we did not expect when we built the list of acts. To better understand what they exactly embrace, we now split the analyse according to these four groups and their content.

#### 2.1. From act 1 to act 10: certification acts

The first group covers different acts such as receiving a manuscript, communicating with authors, finding reviewers and making a review, deciding to accept or not a manuscript, checking information, plagiarism and conflicts of interest. It shows that negative answers are particularly high (ranging from 74% to 90%), and that they peak for the search for reviewers, the evaluation itself, and the acceptance or rejection decision (88%, 90% and 90% respectively). On the other hand, the positive answers range from 9% to 24%.

We labelled this group “certification acts”, as they are involved in the transformation of a manuscript into a valid academic soon-to-be article. The low proportion of monetary transactions is not surprising here. These acts are generally considered as the core of the editorial committee scientific work, which comes in the long tradition of a gift economy in learned societies (Fyfe et al., 2022), as illustrated by the following answer to an open question:

*ID 12754720887: “Editorial board members are acting as volunteers. All software used is free (or licenses are acquired for the entire research institute editing the journal)”*

However, in the few cases where there is a remuneration, it is overwhelmingly attributed to the editor-in-chief. Even if the acts with the highest monetary transactions are at the margins of the scientific evaluation process, they are only partly delegated to external actors that would be paid for checking for plagiarism (money is given to a software/ to a contractor in 12% of journals) or reviewing (referees being remunerated in 2% of journals).

#### 2.2. From act 11 to act 18: material production acts

The second group encompasses act 11 to act 18, including copy editing, language editing, checking compliance with the template, graphic work, proofreading, translation, format production (PDF, HTML), etc. Contrary to the first group, this group shows the highest positive answer rates, distributed between 25% and 44%. The most monetized acts concern the production of the format (pdf, html, xml) (44%), copy editing (37%), language editing (36%), as well as graphic work on the text (35%). These results confirm those of the OADJS study. In the published report that displays a slightly different typology, editing, copy editing, typesetting and design as the most prevalent expenses for Diamond OA journals (Bosman et al., 2021, p. 117-119).

We named this group “material production acts”, acts generally considered as “non-academic work” and the first historically outsourced to printers or publisher. Generally, these acts happen after certification and enable the manuscript to become a digital object within the journal tools. That being said, a majority of the “textual” monetized acts are performed by copyeditors and not contractors. Only format production and translation are mainly subcontracted when monetized. Moreover, the majority of answers were negative (ranging from 54% to 71%), in line with the first point we made about the general trends (figure 1).

#### 2.3. From act 19 to 21: legal dissemination acts

The third group we identified goes from act 19 to 21, including image rights, managing licenses, right management and author contracts. As it is oriented toward the dissemination of the manuscript made article and focus on legal considerations, we named this group “legal dissemination acts”. As for the first group, this one cumulates high rates of negative answers, between 77% and 81%. We might think that the journal prefers to keep the control of legal acts for which its responsibility is at stake, as once again editors in chief and copyeditors represent the majority of beneficiairies when these acts are monetized.

#### 2.4. From act 22 to 26 : technical dissemination acts

The last group (from act 22 to act 26) gathers metadata and DOI assignment as well as online publication of the manuscript into an issue and its associated data. We named this group “technical dissemination acts”, that enable the certified and produced article to circulate from a technical point of view. As far as monetization is concerned, positive answers are higher than in the third group, as we can see for metadata (30%), attribution of a DOI (31%), integration of the text in an issue (30%) and posting online (34%). These acts are still performed by a minority of contractors, but at a much higher rate than legal acts, between 28% and 37% of monetized acts, hence between 8% and 12% of journals.

Technical dissemination acts seem closer to material production acts. As they do not imply scientific or legal responsibility, they are more likely to be monetized. Hence, in the answers to open questions, respondents explain that they pay for some parts of the technical acts (DOI assignment, web dissemination), but never for legal acts.

This section highlighted the actual monetary transactions within Diamond OA journals. Monetary transactions are low for a majority of acts, notably when it comes to scientific evaluation as well as ethical and legal responsibility. Nevertheless, we saw that money – and not only volunteers or direct institutional support – was indeed used to run Diamond OA journals, would it be to pay journal staff or through subcontractors. We now turn to the financial needs of such journals,taking a step further by identifying the acts that could be funded if more money was available.

### 3. Do Diamond OA journals need additional funding and to what end?

If OA Diamond journals already make use of money throughout their publication process, do they need additional fundings? If more money was available, would they be willing to use it? To answer these complex questions, we made use of the notion of “ideal world” in our questionnaire. We define the ideal world as a situation without any financial constraints, where the amount of money available to journals would be unlimited. The ideal world should obviously not be seen as a realistic horizon for the economics of Diamond OA academic publishing. Rather, it is a useful fiction that, as a proxy, enabled us to highlight the acts for which journals would like additional funding or, on the opposite, would not have any use.

Based on this hypothetical situation, we asked Diamond OA journals about their willingness to pay for a given act. The total number of respondents varied, ranging from 202 to 231. To question the willingness to pay, we suggested four possible answers: “yes”, “no”, “maybe”, “I don’t know”. As we can see on the figure 2, the majority of the answers are distributed between “yes” and “no” (the option “maybe” varies between 8% and 16%, while the option “I don’t know” displays even lower proportions, between 3% and 10%).

**Figure 2.**
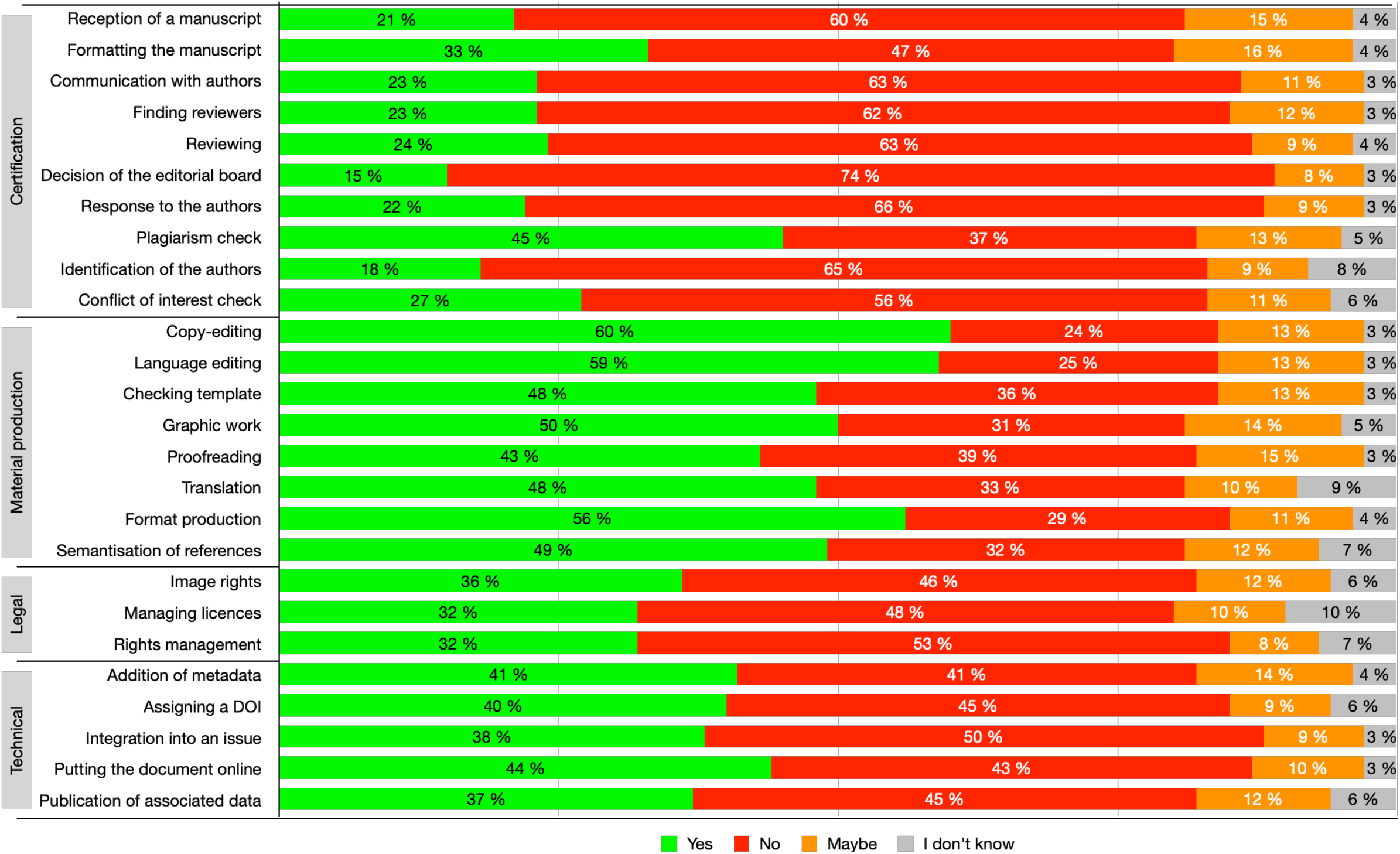
Distribution of willingness to pay in an “ideal world”

In line with the previous figure, answers are organized around the four same groups of acts. This first general result shows that actual monetary transactions and financial needs are well matched. Let us now analyse the answer distribution within each group.

#### 3.1. Funding certification acts in an ideal world

Negative answers are particularly high for certification acts. Journals are generally reluctant to pay for this group of acts, even in a world without financial constraints. As explained by some respondents to the open questions, more money for the certification acts could harm the journal’s scientific independence:

*ID 12765440532* : *« It should be no conflict of interest, and financial support should not limit the action of Editorial board and editor in chief in selecting the content of the journal »*.

As a consequence, a respondent asks for editorial independence certainty in the eventuality of direct funding mechanism:

*ID 12798245985: « If it were funded externally in any way, the funding would have to be offered with few (or no) strings attached. The journal needs to be independent to ensure academic freedom »*.

Hence, the decision to accept or reject a manuscript, which is the scientific decision *par excellence*, reaches the highest value with 74% of “No”. The following acts which are still an important part of the editorial committee’s work (communication to authors, finding reviewers and reviewing, answering to and identifying authors) have also high negative response rates, between 59% and 66%.

Despite the massive negative tendency, we notice one exception. The act “plagiarism check” is the only one for which positive answers are superior to negative ones (37% “no” versus 45% “yes”). One possible explanation is that plagiarism check is mostly considered at the margin of the editorial committee’s scientific work, routinely achieved by the use of a software, be it internal or external of the journal.

#### 3.2. Funding the material production in an ideal world

We previously saw that monetary transactions had more chances to occur on material production acts. In an ideal world without financial constraints, this trend is massively confirmed. Not only material production acts gather the highest positive answers, but the positive answers are also largely superior to the negative ones. In a world free of financial constraints, the acts for which journals would be most willing to pay are copy editing (60%), language editing (59%) and format production (56%). The remaining answers ranged from 43% to 50% for compliance checking, graphic work, proofreading, translation, and semantization of references^4^.

Hence, figure 2 gives a clear message: as far as material production acts are concerned, Diamond OA journals would pay for it if they had access to external funds in addition to their self-support ecosystem. According to some of our respondents, financial capacities to outsource material production acts could be an opportunity to focus on the core of editorial tasks, what we called “certification acts”:

*ID 12772112368: “In an ideal world, we would delegate a lot of the jobs that are related to design, production and copy editing, in order to focus on the management of the editorial process and editorial selection / management*.

*ID 12797718435* : *“In an ideal situation, all formal and technical tasks related to the publishing processes would be out-sourced in order to help the editors to focus on organizing peer review and communicating with authors”*.

#### 3.3. Funding legal dissemination acts in an ideal world

Legal acts of dissemination concentrate high negative answers, even if they are still inferior to negative answers of the first group. Journals tend to not embrace a monetary transaction for image rights (46%), license management (48%) and author contracts (53%). Even if they had access to unlimited funds, journals editorial teams still put at stake their legal responsibility when it comes to those acts. These results confirm what we saw insection 2 on actual Diamond journals’ monetary transactions.

#### 3.4. Funding technical dissemination acts in an ideal world

The trend for technical acts of dissemination is more nuanced than the previous group. Negative answers prevail, except for addition of metadata (“yes” and “no” reach both 41%) and online publication (positive answers hardly prevail with a rate of 44%). Nonetheless, Diamond OA journals are more willing to make use of money for technical dissemination acts than legal ones.

From the ideal world perspective, the answers for the two first groups of acts delivered clear-cut messages. Funds are not welcome to support certification acts (with the exception of plagiarism check), while they seem decisive in many cases for material production acts. By comparison, the answers for the dissemination acts are fuzzier. Negative answers are globally higher than the positive ones, but it seems that an important proportion of journals could make use of funds, notably to ease the achievement of technical acts of dissemination.

Another way to look at these results is to compare the current situation with the ideal situation, to isolate the effect of unlimited funding. For all the tasks, we considered the changes that a new flow of funds would bring about among those journals that do not currently monetise them. All tasks would be subject to additional monetisation, in proportions ranging from 14% to 52% of these respondents. If we shift from tasks to journals, we find that half of the latter would make use of supplementary money for at least four tasks, while 59% would need it for at least one task.

If a direct funding mechanism would be useful for a majority of journals, a majority of tasks would not be affected by financial flows. This result shows both the importance of institutional and volunteer support, and the non-profit nature of Diamond OA journals. To our own question on their financial goals, only 4 journals answered they were supposed to make profit, in line with the OADJS study. Hence, in the current Diamond OA ecosystem, a new funding stream would quasi exclusively be spent to support publication, rather than being redistributed to owners and shareholders. That is, under the condition journal would be able to receive and spend money.

### 4. Are Diamond OA journals able to engage in monetary transactions?

After having shown that Diamond OA journals do need extra money, we now investigate a second, related stake: the journal’s ability to manage monetary transactions. At first sight, addressing such a question seems simple ; nevertheless, as we built the questionnaire, we realized that it was trickier than it seems.

First, on the definition of “monetary transactions”. One could think that these terms only refer to the fact of receiving funds, that is, a money transfer from a research funder to a Diamond OA journal. Nevertheless, to be fully operational, a direct funding mechanism relies on the journals’ ability not only to receive, but also to possibly spend money to buy some services that are not directly supported in its ecosystem. The parts of the questionnaire related to monetary transactions clearly distinguish between the receiving capacity and the spending capacity. Hence, in what follows, we study the journals’ capabilities to receive money, to spend money, and to do both.

Second, we consider that the notion of “ability” to receive or spend money may refer to three different situations. In the more obvious one, journals do currently receive money and make spendings. Alternatively, journals that are not able to make direct transactions, may perform it via an intermediary that would act on its behalf. It is notably the case for certain small journals which do not have a legal capacity to deal with money but rely on a dissemination platform or other third party to manage their funds. Given that the Diamond ecosystem is characterized by its small-scale economy, forgetting that second situation would lead to overlook a part of the reality of Diamond OA journals’ transactional capacities. Furthermore, the OADJS data for our responding journals confirms this small-scale economy, with 85% of them paying 0 to 2 full-time equivalent persons for operational work, and a third of the journals paying less than $1,000 for the same work. To distinguish those two situations, we decided to call the first one “direct transaction capacity” and the second one “mediated transaction capacity”. Additionally, if a journal does not have neither direct nor mediated capacity, it may develop one in the future, with the perspective of a recurrent direct funding mechanism. To address this third situation, we coined the category “conditional transaction capacity”, that is, the capacity to develop a transaction capacity as a counterpart for sufficient funding from research funders.

To understand the journals’ capacities to engage into monetary transactions, we successively focus on these three categories: direct transaction capacity, intermediate transaction capacity, and conditional transaction capacity.

We first asked journals about direct transaction capacity. The major result is that a very large majority of them can receive or spend money among the 254 journal answers (70% on the receiving side, and 73% on the spending side). Nevertheless, only 151 respondents (60%) can both receive and spend money at the same time. All the respondents who didn’t answer positively to the questions about direct transaction capacities were asked about mediated transaction capacities. We obtained 72 answers regarding mediated transaction capacities, with respectively 35% and 22% positive answers, but only 12 respondents (17%) can both receive and spend money through an intermediary. This asymmetry between spending and receiving confirms our methodological precaution that leads us to distinguish between the two sides. We shall add that an approximate quarter of respondants which state that they do not know about mediated transaction capacities (24% for the receiving side vs. 28% for the spending side), which indicates that these schemes are often not envisioned in the current economy of these journals.

As far as conditional transaction capacities are concerned, we did not distinguish here between the two sides of the transaction. We considered that, if a journal acquires transaction capacities with the specific purpose of managing research funder’s money, it will settle a system that allows to manage money reception and expense alike. We asked the journals which did not have neither direct nor mediated transaction capacity if they would develop such a transaction system. Among the 50 answers, a majority were negative: 42% of the respondents do not have what we call a conditional transaction capacity. Nevertheless, 30% of them would develop such a transaction system, while more than a quarter (28%) are still undecided. Since the size of the sample is quite small, the results regarding the willingness of journals without a transaction system to adopt one must be taken with caution. However, the positive answers (n=15 journals, 30%) can be considered as a convincing trend about the declared capacity of these journals to organize themselves to receive and spend money or, conversely, to support a capacity centre to delegate these transactional abilities as recommended by the OADJS (Becerril et al., 2021).

In order to gain a more comprehensive view of Diamond OA journals capacities, it is possible to sum the conditional capacity with the direct and mediated one so as to form what we call a “total transaction capacity”. In the table 2 below, we summarized the data on transaction capacities already shown with a distinction between the receiving side, the spending side, and the ability to do both. We also added two pieces of information: the total transaction capacity in absolute value and in percentage.

**Table 2.** Total transaction capacities (n=254)

| **Types of transaction capacity** | **Receiving side** | **Spending side** | **Receiving and spending** |
| --- | --- | --- | --- |
| Direct transaction capacity | 178 | 185 | 151 |
| Mediated transaction capacity | 25 | 16 | 12 |
| Conditional transaction capacity | 15 | 15 | 15 |
| **Total transaction capacity (N)** | **218** | **216** | **178** |
| **Total transaction capacity (%)** | **86%** | **85%** | **70%** |

While the previous section clearly showed that Diamond OA journals had monetary needs, this section highlights a crucial result: for 6 out of 7 Diamond OA journals can either receive or spend money, and 7 out of 10 Diamond OA journals can already manage both, or could engage into such activites under the condition of sufficient funds from RFOs. This information does not invalidate the need for a “capacity centre” as recommended by OADJS, but a direct funding mechanism would not need the establishment of such a device to already be very useful for a majority of Diamond OA journals.

### 5. Meeting RFOs requirements

We finally anticipated the expectancies of RFOs involved in this new financial stream, based on what they usually ask researchers who have received funding. That is, to acknowledge the funding by indicating the name of the funder and the grant number in their research outputs. In fact, RFOs are usually neither authors nor cited, but are part of the things, people and institutions that have been needed, hence named somewhere into an acknowledgement section which rewards them for their monetary contribution (Cronin and Weaver, 1995). For the past fifteen years, a specialized literature has dealt with these funding acknowledgements and has regularly shown that the information they contain is often incomplete despite mandates issued by RFOs within their grants (for a review, see Alvarez-Bornstein and Montesi, 2020). Faced with this, more and more actors are campaigning for the use of permanent identifiers for RFOs in order to simplify and standardize funder metadata, particularly because of their open access policies which lead them to have not only funded the research, but also the publication itself. So, in a publishing world more and more revolving around transparency and openness principles, we considered the development of new funding channels as demanding the same requirements that already exist for APC and subscription journals.

This section examines the propensity of journals to ensure the traceability of research funders. The first part highlights the current capacities of journals to provide visibility to research funders (5.1.) As Diamond funding mechanisms have to be distinguished from the APC model, we asked journals about their ability not only to match funding metadata, but to report on a yearly basis for any given RFO. The second part focuses on journals that currently lack reporting capacity, and develops thoughts on potential incentives for journals (5.2).

#### 5.1. Making the funders’ contribution visible

Making visible the contribution of funders starts at the level of individual articles: the challenge is to be able to match, for a given article, on which funding basis the authors carried out their research and produced their manuscript. This match clearly rests on the authors side, hence our question to journals: “For each article you publish, do you ask authors to indicate the RFO of the research or grant identification that led to its publication?”. Among the 232 answers, 56% of the journals declare that they carry out minimal acknowledgment of research funding. Symmetrically, a significant proportion of journals (42%) do not perform any such acknowledgment, while only 1% of respondents were unable to answer. That could seem a rather low rate, but the literature shows that less than half of the Web of Science journal articles were showing funding metadata in 2016 (Tang et al., 2017).

While the acknowledgment of funding by article is key to any funding information, in order to achieve complete visibility, it would be necessary for journals to produce a report that lists RFOs and aggregates the articles that they have contributed to produce. When asked about that, journals answers show a reversed trend: of the 136 respondents, only 36% claim to be currently able to produce such a report, while 40% consider they could not. We notice a large proportion of respondents who simply do not know whether reporting to funders is possible (24%). If we add the fact that 96 journals didn’t answer this question while they did to the previous one, we have unveiled a new piece of evidence of the lack of RFOs presence in the Diamond ecosystem.

#### 5.2. Incentives for the development of a reporting system

The previous sections indicated that Diamond OA journals were ready to welcome money from research funders for two reasons: they need this money to achieve certain acts of the publication process, and they are for a large part able to receive and spend it. This section on the funders’ visibility requirements highlights a possible pitfall for the implementation of direct funding mechanisms. As we saw above, an important part of Diamond OA journals does not currently acknowledge funding information for each article. Moreover, this part grows when it comes to produce a systematic reporting. Obviously, the current situation regarding funders’ requirements derives from the fact that very few funders currently foster Diamond OA journals or even the research that gets published in these outputs. Hence, we asked to what extent journals would develop a reporting system as a counterpart for direct funds.

To further explore the conditions under which journals would be able to acquire technical reporting capacities, we addressed the question of potential incentives: “If research funding organizations required this type of report in order to provide regular funding to your journal, would you be likely to put it in place?”. The hypothesis of a regular income seems to largely increase the willingness to take charge of reporting practices. 73% of the 175 respondents which have not the current reporting capacity would agree to adopt a reporting scheme. Negative answers were extremely low (7% of the population). We also note that the proportion of undecided remains low, with one fifth of journals (19%).

With a hypothetical incentive in place, what level of income would be sufficient for journals to adopt such a reporting system? We analysed the few qualitative answers obtained on this question (8 in total). First of all, these journals cover a wide variety of disciplines, ranging from mathematics to computer sciences, biology, history, literature, law and linguistics; but also a variety of countries (France, South Africa, Italy, Australia, as well as several international journals). Among the answers mentioning them, the annual monetary amounts vary widely: from $2,000 for the minimum threshold to a range between $8,000 and $20,000 for the majority of answers, that is the equivalent of a few APCs. These amounts have a related condition: not creating additional work and costs to the publication process. For example, some journals advocate for a technical support to make it easy to produce. The possibility of hiring someone to write the report, such as a copy editor, is also a strongly considered option.

Despite the eventuality of obtaining a regular income for the journal, a small minority of respondents declined the setting of reporting (n=13 journals in total, or 7% of responses). Nine of them agreed to provide explanations in response to the open-ended question. Let us emphasize once again the diversity of disciplines (postcolonial studies, social sciences, geography, biology, materials science, etc.) and countries that make up this sub-sample (France, Italy, USA, and several international journals, including a group of Middle Eastern countries). Two main arguments are put forward to explain the refusal to set up a reporting system. First, some journals explain they do not need additional funding, considering that they rarely deal with RFOs, or that the research behind the submitted manuscripts was not granted by RFOs. Second, the idea of reporting is not desirable for some journals, because of the additional administrative work it would entail and the risk of losing the journal’s independence.

## Conclusion

We started this article by insisting on the political and ethical problems that APCs entail, and the urge to develop a sustainable Diamond open access world. To contribute to this perspective and give practical tools to go beyond APCs, we followed one OADJS’ key recommendation: we examine the practical conditions to set up a direct funding for Diamond OA journals. We have drawn four main threads that allow us to understand the practical modalities of such direct funding.

We examined first the monetary needs of Diamond OA journals all along the publication process. We saw that a vast majority of Diamond OA journals already make use of money to produce scientific papers for certain acts, notably concerning the material production of the article and the technical acts of its dissemination. Second, we demonstrated that those needs would be fulfilled in the eventuality of a regular income from research funders, lowering the volunteers’ burden. Third, we tackled the question of Diamond journal’s capacity to manage monetary transactions. A vast majority of them can receive and spend money or could be able to do so in the eventuality of a direct funding mechanism. Lastly, we looked at the funders’ requirements. If a small majority of Diamond OA journals currently have funding acknowledgments for each paper, it is still far from being met for lots of respondents. Nevertheless, we saw that this point could be enhanced with the promise of regular income through a direct funding mechanism.

A logical follow-up of our work would be to inquire RFOs about the feasibility of the direct funding on their side. Even if the national Austrian funding agency has done it in the past (Rieck, 2019), it is in particular, their legal ability to directly subsidize journals that should be questioned. If RFOs main scheme is to support research projects or institutions through grants, journal funding is far less common. Then, the empirical scheme of a direct funding mechanism should be treated. As a starting point, we can suggest four settings that are based on existing models in the open access world, whether Diamond or not.

The first one would be a matter of defining a “journals list” that would receive support simply by virtue of their title being part of it. For example, such a list could be built through the combined criteria of “DOAJ inclusion”, “license type”, “journal country” or in considering mission-value criteria. A second setting could include a minimum usage criterion to deliver money. For example, for a given RFO, this would be journals that have published at least one article from a research project that has received funding from them in the last two years. Such a system has been set up by some universities, notably the University of Amsterdam, for Diamond OA journals that would have published articles from one of their academic staff. A third setting is inspired by the publisher ACM and its tier-based “transformative agreement” model (Dufour et al., 2021). This model divides publication volumes into 10 different tiers. The highest tier gave access for an unlimited number of published articles. The lowest allowed for 1 to 3 articles per year. Each intermediary tier has a fixed fee for a certain volume of publication. Using this model for Diamond OA journals, the highest tier would be the maximum amount of support from a given RFO, given above a certain level of reported articles. Finally, a fourth system would be an APC-like aggregated system for Diamond OA journals, where the journal sums up the number of articles of interest for a funder and receives in return an amount that depends linearly on this number of articles. Here, the link between money flows and volumes of publications is comparable to an APC model; nonetheless, no author nor reader would pay. All these schemes would have in common to be recurrent ones, building a constant monetary flow from RFOs to journals.

If Diamond OA journals are globally ready and eager to welcome direct funding mechanisms, the permanent monetary flow induces new challenges. The first one is administrative, given that an accounting and reporting system can imply a new administrative task for the editorial team. The second challenge is a well-known ethical one. Indeed, financial dependency could harm the independence and integrity of the editorial policy: an incentive to publish “funding” authors (that is, the ones supported by a research funder), or, as a more general rule, the laxing of peer review to enhance funding sources. However, this threat is not specific to Diamond OA journals since it has been suspected at observed at some APC OA publishers. Third, in line with the predatory journals that have developed as parasites of the APC model, several of our respondents warned of the possible emergence of Diamond predatory journals. Even without going that far, the question of the increase, displacement and decrease of inequalities specific to the academic world will have to be an integral part of the evaluations of direct funding mechanisms. Only then can this new financial stream be deemed sustainable for Diamond OA journals.

## Acknowledgments

This work was funded by the French Ministry of Higher Education, Research and Innovation through the French Open Science Committee. We would like to thank Odile Contat, Marin Dacos and Claire Leymonerie as Ministry advisers for the study, Victoria Brun, Morgan Meyer and our colleagues at CSI for their insightful comments, our colleagues of the French Open Science Committee and our colleagues of the DIAMAS project who have made constructive remarks on earlier versions of this text.

## Appendix 1 Introduction page of the questionnaire

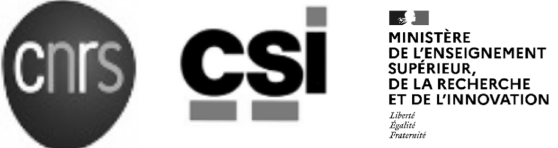

#### New Funding Mechanisms for OA Diamond Journals

##### Introduction

**You answered the <u>OA Diamond Journal Survey</u> led by the <u>OPERAS consortium</u> in 2020. You agreed to be contacted for the follow-ups of this study, and we thank you for that. You are invited to participate in a complementary survey dedicated to test the feasibility and desirability of developing funding mechanisms for OA Diamond journals. In line with one of the <u>OA Diamond Study Report recommendations</u>, we would like to identify practical financial schemes where institutional research funders could finance OA Diamond Journals**.

##### Goal

**This study, complementary to the one conducted in 2020, aims to test the feasibility and desirability of such funding mechanisms. The questionnaire we are sending you will help us to identify OA Diamond journals’ financial needs, their ability to report, their opinion and comments on such funding schemes. The results of this study will be gathered to explore different Diamond funding schemes with research funding organisation**.

##### Context of the study

**Your answers and those of many other journals enabled OPERAS’ members to publish outstanding results and recommendations in the recently released OA Diamond Journal study report. Amongst the many support measures envisaged for the whole Diamond ecosystem, the report encourages the development of funding mechanisms specifically directed to Diamond journals. One way of achieving this goal is to lean on already existing research funding organizations, that could reallocate part of their monetary flows from APCs to original diamond funding mechanisms**.

##### Instructions

**The questionnaire is organized around five sections :**

1. **Your journal identification**
2. **Your journal economic configuration**
3. **Tasks on a given manuscript**
4. **Funding and grant reports**
5. **Your opinion on funding mechanisms**

**You can download the complete list of questions in a PDF file before providing your answers. As a follow-up survey, this questionnaire is much shorter than the one led by OPERAS in 2020. It should take you less than 30 minutes to complete. The questionnaire is open until July 8, 2021**.

##### Team and funder

**We are researchers at the <u>Center for the Sociology of Innovation</u> (Mines ParisTech and CNRS). Our work is financed by the <u>Committee for Open Science</u> (CoSo) as part of** the **French Ministry for Higher Education and Research. Within the OPERAS consortium, we participated in the 2020 OA Diamond Journal Survey**.

##### Contacts and questions

**Thank you for your contribution and don’t hesitate to contact us if you have any question or issue answering the survey. You can reach us at the following addresses :**

## Appendix 2 Questionnaire Content

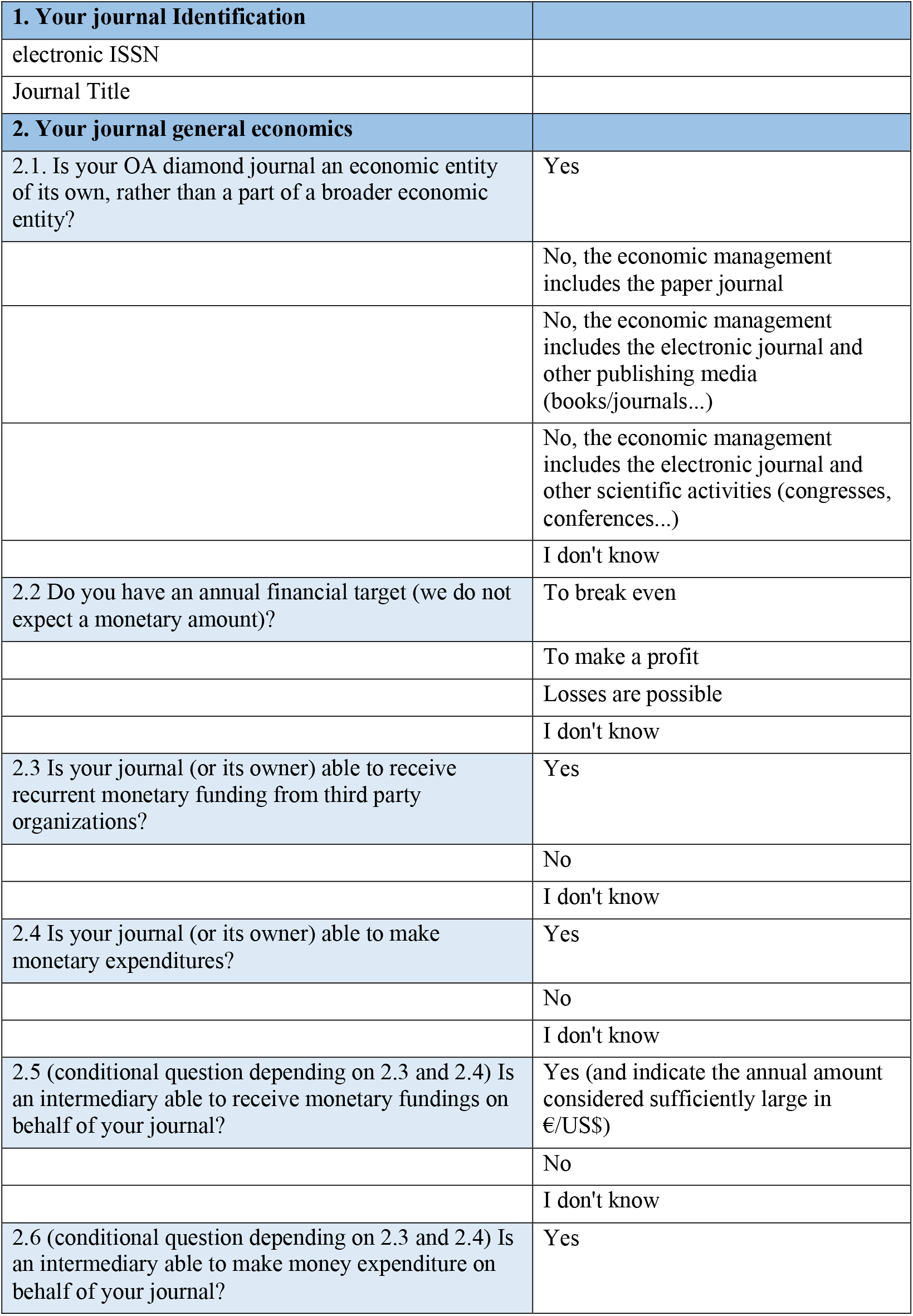
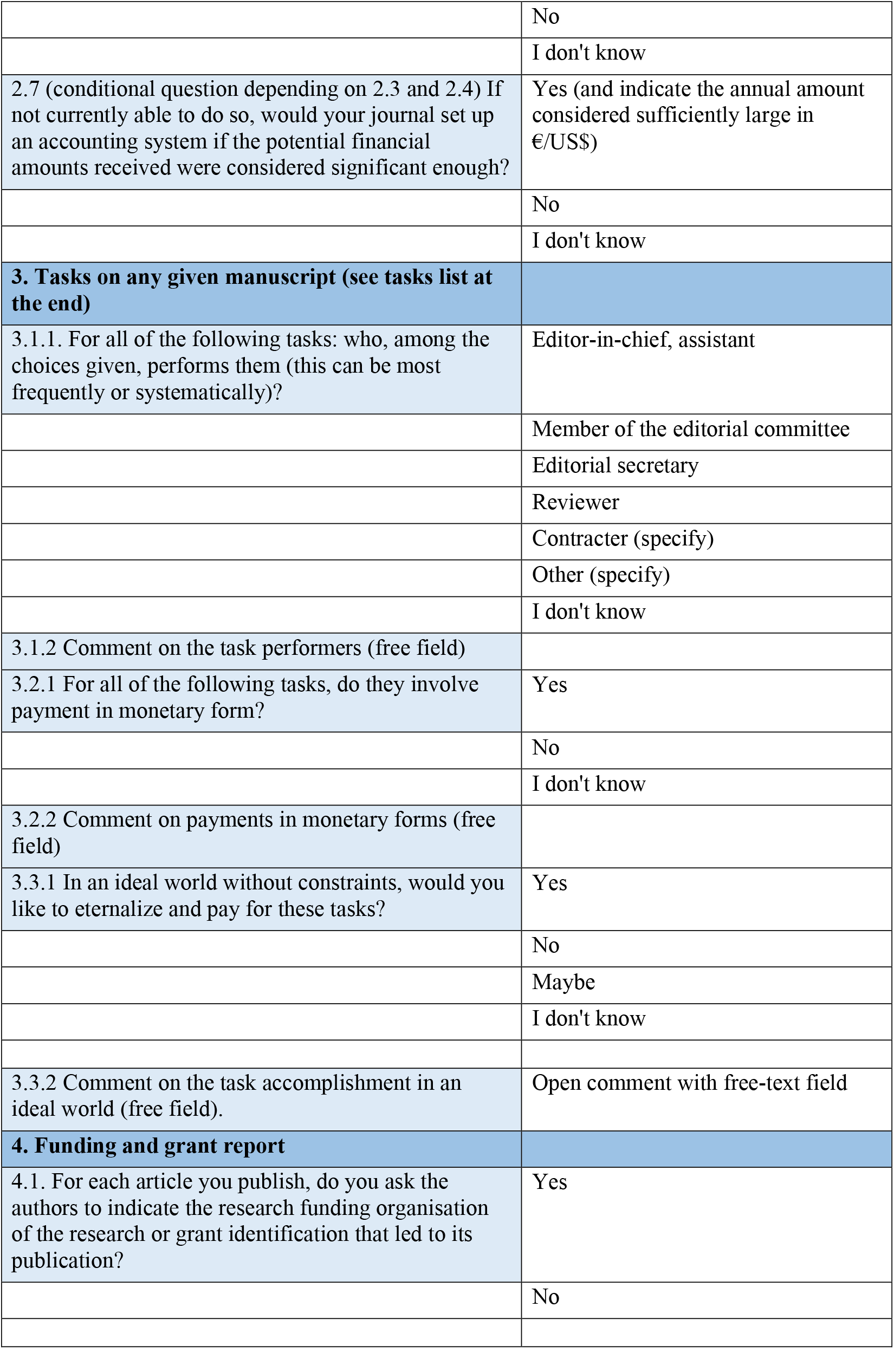
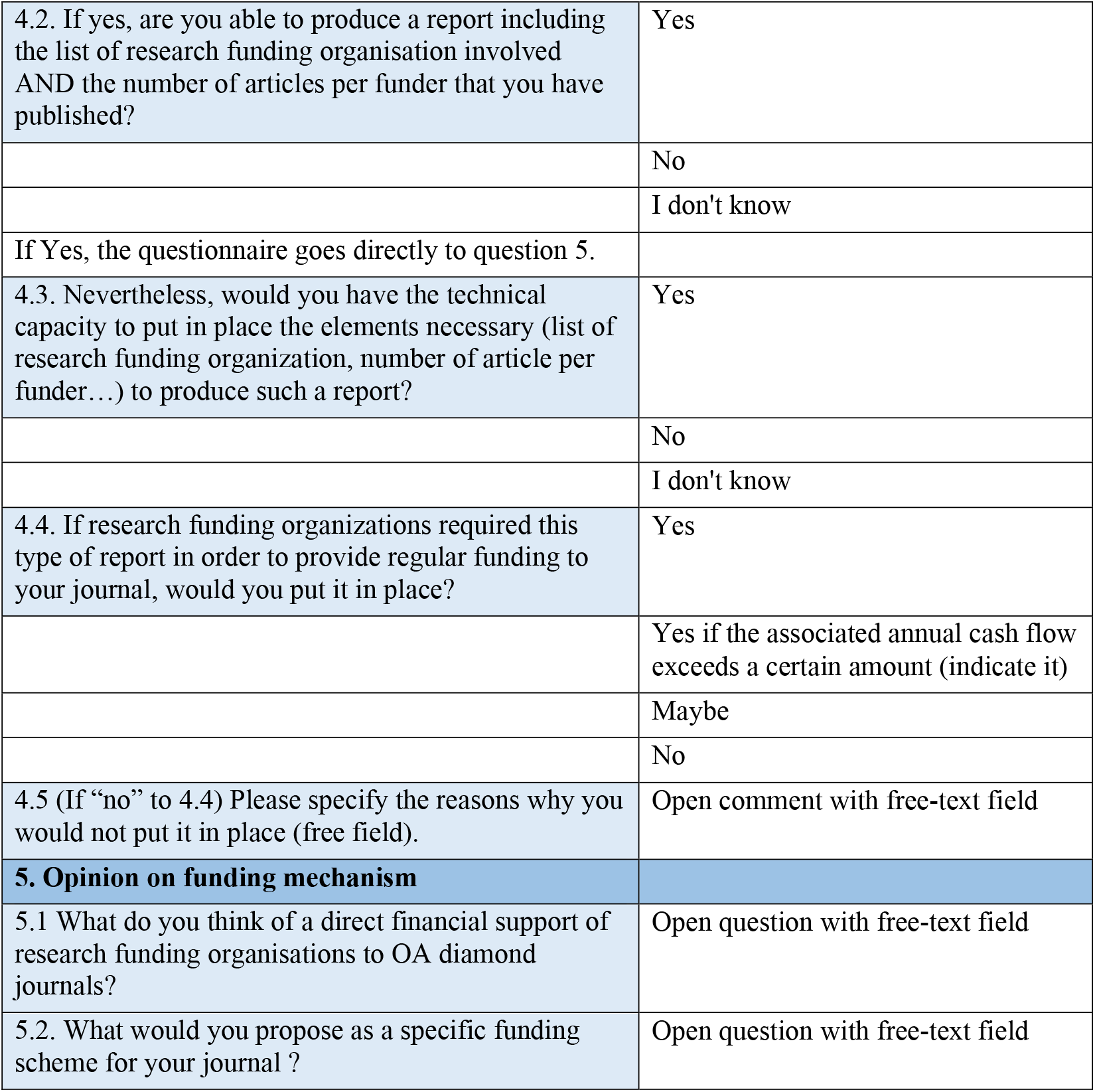

Appendix 2 – Questionnaire Content
|  |  |
| --- | --- |
| **1. Your journal Identification** |  |
| electronic ISSN |  |
| Journal Title |  |
| **2. Your journal general economics** |  |
| 2.1. Is your OA diamond journal an economic entity of its own, rather than a part of a broader economic entity? | Yes |
|  | No, the economic management includes the paper journal |
|  | No, the economic management includes the electronic journal and other publishing media (books/journals...) |
|  | No, the economic management includes the electronic journal and other scientific activities (congresses, conferences...) |
|  | I don't know |
| 2.2 Do you have an annual financial target (we do not expect a monetary amount)? | To break even |
|  | To make a profit |
|  | Losses are possible |
|  | I don't know |
| 2.3 Is your journal (or its owner) able to receive recurrent monetary funding from third party organizations? | Yes |
|  | No |
|  | I don't know |
| 2.4 Is your journal (or its owner) able to make monetary expenditures? | Yes |
|  | No |
|  | I don't know |
| 2.5 (conditional question depending on 2.3 and 2.4) Is an intermediary able to receive monetary fundings on behalf of your journal? | Yes (and indicate the annual amount considered sufficiently large in €/US\$) |
|  | No |
|  | I don't know |
| 2.6 (conditional question depending on 2.3 and 2.4) Is an intermediary able to make money expenditure on behalf of your journal? | Yes |
|  | No |
|  | I don't know |
| 2.7 (conditional question depending on 2.3 and 2.4) If not currently able to do so, would your journal set up an accounting system if the potential financial amounts received were considered significant enough? | Yes (and indicate the annual amount considered sufficiently large in €/US\$) |
|  | No |
|  | I don't know |
| **3. Tasks on any given manuscript (see tasks list at the end)** |  |
| 3.1.1. For all of the following tasks: who, among the choices given, performs them (this can be most frequently or systematically)? | Editor-in-chief, assistant |
|  | Member of the editorial committee |
|  | Editorial secretary |
|  | Reviewer |
|  | Contracter (specify) |
|  | Other (specify) |
|  | I don't know |
| 3.1.2 Comment on the task performers (free field) |  |
| 3.2.1 For all of the following tasks, do they involve payment in monetary form? | Yes |
|  | No |
|  | I don't know |
| 3.2.2 Comment on payments in monetary forms (free field) |  |
| 3.3.1 In an ideal world without constraints, would you like to eternalize and pay for these tasks? | Yes |
|  | No |
|  | Maybe |
|  | I don't know |
| 3.3.2 Comment on the task accomplishment in an ideal world (free field). | Open comment with free-text field |
| **4. Funding and grant report** |  |
| 4.1. For each article you publish, do you ask the authors to indicate the research funding organisation of the research or grant identification that led to its publication? | Yes |
|  | No |
| 4.2. If yes, are you able to produce a report including the list of research funding organisation involved AND the number of articles per funder that you have published? | Yes |
|  | No |
|  | I don't know |
| If Yes, the questionnaire goes directly to question 5. |  |
| 4.3. Nevertheless, would you have the technical capacity to put in place the elements necessary (list of research funding organization, number of article per funder...) to produce such a report? | Yes |
|  | No |
|  | I don't know |
| 4.4. If research funding organizations required this type of report in order to provide regular funding to your journal, would you put it in place? | Yes |
|  | Yes if the associated annual cash flow exceeds a certain amount (indicate it) |
|  | Maybe |
|  | No |
| 4.5 (If “no” to 4.4) Please specify the reasons why you would not put it in place (free field). | Open comment with free-text field |
| **5. Opinion on funding mechanism** |  |
| 5.1 What do you think of a direct financial support of research funding organisations to OA diamond journals? | Open question with free-text field |
| 5.2. What would you propose as a specific funding scheme for your journal ? | Open question with free-text field |

## Footnotes

1 Of course, some initiatives make use of funders to support Diamond OA journals, but it has a limited impact on the Diamond ecosystem. This limitation is notably due to their attachment to an institution, a discipline, or at best to a national institution, without global coordination, despite projects such as The Global Sustainability Coalition for Open Science Services (SCOSS).

2 Secondary exploitation of qualitative data from OADJS, not shared due to the inability to guarantee anonymization of the data.

3 The 7 others have probably been informed of the survey by one of the primary journals. Their answers were taken into account in the following results.

4 Semantization of references is the process by which each reference at the end of an academic article is associated with its DOI, and become clickable for a reader on a digital device.

## References

Ancion, Z., Borrell-Damián, L., Mounier, P., Rooryck, J., Saenen, B., 2022. Action Plan for Diamond Open Access. https://doi.org/10.5281/zenodo.6282403

Anderson, K., 2018. Focusing on Value — 102 Things Journal Publishers Do (2018 Update) The Scholarly Kitchen. URL https://scholarlykitchen.sspnet.org/2018/02/06/focusing-value-102-things-journal-publishers-2018-update/ x(accessed 3.17.23).

Becerril, Arianna, Bosman, Jeroen, Bjørnshauge, Lars, Frantsvåg, Jan Erik, Kramer, Bianca, Langlais, Pierre-Carl, Mounier, Pierre, Proudman, Vanessa, Redhead, Claire, Torny, Didier, 2021. OA Diamond Journals Study. Part 2: Recommendations. Zenodo. https://doi.org/10.5281/ZENODO.4562790

Bosman, J., Frantsvåg, J.E., Kramer, B., Langlais, P.-C., Proudman, V., 2021. OA Diamond Journals Study. Part 1: Findings. Zenodo. https://doi.org/10.5281/zenodo.4558704

Contat, O., Gremillet, A.-S., 2015. Publier : à quel prix ? Étude sur la structuration des coûts de publication pour les revues françaises en SHS. rfsic. https://doi.org/10.4000/rfsic.1716

Crow, R., Gallagher, R., Naim, K., 2020. Subscribe to Open: A practical approach for converting subscription journals to open access. Learned Publishing 33, 181–185. https://doi.org/10.1002/leap.1262

Dufour, Q., Pontille, D., Torny, D., 2021. Contracter à l’heure de la publication en accès ouvert. Une analyse systématique des accords transformants (report). CNRS ; Comité pour la science ouverte. https://doi.org/10.52949/2

Ellers, J., Crowther, T.W., Harvey, J.A., 2017. Gold Open Access Publishing in Mega-Journals: Developing Countries Pay the Price of Western Premium Academic Output. Journal of Scholarly Publishing 49, 89–102. https://doi.org/10.3138/jsp.49.1.89

Estelle, L., Wise, A., 2023. OASPA Equity in Open Access Workshop 1 Report. Zenodo. https://doi.org/10.5281/zenodo.7733869

Fyfe, A., Moxham, N., McDougall-Waters, J., Mork Rostvik, C., 2022. A History of Scientific Journals. Publishing at the Royal Society, 1665–2015. UCL Press.

Grossmann, A., Brembs, B., 2021. Current market rates for scholarly publishing services. F1000Res 10, 20. https://doi.org/10.12688/f1000research.27468.1

Khoo, S.Y.-S., 2019. Article Processing Charge Hyperinflation and Price Insensitivity: An Open Access Sequel to the Serials Crisis. LIBER 29, 1–18. https://doi.org/10.18352/lq.10280

Laakso, M., Solomon, D., Björk, B.-C., 2016. How subscription-based scholarly journals can convert to open access: A review of approaches. Learned Publishing 29, 259–269. https://doi.org/10.1002/leap.1056

Miedema, Frank, Verbeke, Demmy, Sondervan, Jeroen, Woutersen-Windhouwer, Saskia, Oort, Frans, 2020. Beyond APC: On the Need for Diamond Open Access Publication Platforms. https://doi.org/10.5281/ZENODO.4758334

Morrison, H., Borges, L., Zhao, X., Kakou, T.L., Shanbhoug, A.N., 2022. Change and growth in open access journal publishing and charging trends 2011–2021. Journal of the Association for Information Science and Technology 73, 1793–1805. https://doi.org/10.1002/asi.24717

Mounier, P., 2012. Freemium as a sustainable economic model for open access electronic publishing in humanities and social sciences1. ISU 31, 225–233. https://doi.org/10.3233/ISU-2012-0652

Pooley, J., 2021. Collective Funding to Reclaim Scholarly Publishing. Commonplace. https://doi.org/10.21428/6ffd8432.250139da

Rieck, K., 2019. The FWF’s Open Access Policy over the last 15 Years –Developments and Outlook. Mitteilungen der VÖB 72, 408–423. https://doi.org/10.31263/voebm.v72i2.2837

Ross-Hellauer, T., Reichmann, S., Cole, N.L., Fessl, A., Klebel, T., Pontika, N., 2022. Dynamics of cumulative advantage and threats to equity in open science: a scoping review. R. Soc. open sci. 9, 211032. https://doi.org/10.1098/rsos.211032

Smith, A.C., Merz, L., Borden, J.B., Gulick, C.K., Kshirsagar, A.R., Bruna, E.M., 2021. Assessing the effect of article processing charges on the geographic diversity of authors using Elsevier’s “Mirror Journal” system. Quantitative Science Studies 2, 1123–1143. https://doi.org/10.1162/qss_a_00157

Solomon, D.J., Björk, B.-C., 2012. Publication fees in open access publishing: Sources of funding and factors influencing choice of journal. Journal of the American Society for Information Science and Technology 63, 98–107. https://doi.org/10.1002/asi.21660

Suber, P., 2007. Flipping a journal to open access. SPARC Open Access Newsletter.

Tang, L., Hu, G., Liu, W., 2017. Funding acknowledgment analysis: Queries and caveats. Journal of the Association for Information Science and Technology 68, 790–794. https://doi.org/10.1002/asi.23713

The Budapest Open Access Initiative:20th Anniversary Recommendations 2022. URL https://www.budapestopenaccessinitiative.org/boai20/ x(accessed 3.28.23).

Verbeke, D., Mesotten, L., 2022. Library funding for open access at KU Leuven. Insights 35, 1. https://doi.org/10.1629/uksg.565

Waidlein, N., Wrzesinski, M., Dubois, F., Katzenbach, C., 2021. Working with budget and funding options to make open access journals sustainable. Zenodo. https://doi.org/10.5281/zenodo.4558790

Wise, A., Estelle, L., 2020. How society publishers can accelerate their transition to open access and align with Plan S. Learned Publishing 33, 14–27.

Wise, A., Estelle, L., 2019. How libraries can support society publishers to accelerate their transition to full and immediate OA and Plan S. Insights 32.

Yang, A., Asa, N., Krause, A., Stewart, B., Farley, A., 2023. Policy opportunities: Economics of academic publishing. https://doi.org/10.21955/GATESOPENRES.1117017.1

